# A new paradoxosomatid millipede, *Manikidesmus suriensis* (Polydesmida: Paradoxosomatidae) from West Bengal, India

**DOI:** 10.1101/2021.04.25.441377

**Authors:** Somnath Bhakat

## Abstract

A new genus and species of the family Paradoxosomatidae from southern part of West Bengal, India is described. The new genus *Manikidesmus* gen. n. is diagnosed by combination of following characters: reduced paranota, distinct pleural keel, unpaired sternal lamella on 5^th^ sternite, prefemur with setal brush, setal brush on tibia and tarsus in male, lamina medialis long straight with a curved hook, expanded post femoral lamina with a spine and tibiotarsus with a spine on the distofemoral process.

The genus is distinguished from its Indian congeners by one or more diagnosed characters. In *Manikidesmus suriensis*, sp. n. tibia and tarsus bear setal brush in male (vs. absent in *Oxidus* and *Chondromorpha*), gonopod femur short, flat and without post femoral demarcation (vs. long, thin cylindrical with post femoral demarcation in *Polydrepanum* and *Orthomorpha*), tibiotarsus of gonopod long and a spine on the distofemoral process (vs. short and without spine in *Anoplodesmus*). In *Kronopolites*, coxa of gonopod densely bristled, collum with two rows of long bristles, femur long, slender and with a spine. All these characters are absent in the present genus. In *Streptogonopus*, gonopod femorite is distinctly demarcated from post femorite region and solenomere is twisted with solenophore but in the present species, femorite is not be separated from post femorite region and solenomere is free from solenophore.

## Introduction

Paradoxosomatidae is one of the largest families in the entire class of Diplopoda, dominating the millipede fauna of Indo-Australia and currently comprised of 200 genera and more than 1000 species (Jeekel, 1968; Nguyen and Sierwald, 2013; Likhitrukarn et al., 2014). Paradoxosomatidae has three subfamilies viz. Alogolyxinae, Australiosomatinae and Paradoxosomatinae. Most of the species belongs to Paradoxosomatinae (about 760 species). In India, the only tribe Polydrepanini of the subfamily Alogolyxinae and six tribes Nedyopodini, Orthomorphini, Paradoxosomatini, Sulciferini, Sundaninini, and Tectoporini of the subfamily Paradoxosomatinae are available (Nguyen and Sierwald, 2013). Golovatch and Wesener (2016) reported 22 genera and 56 species of Paradoxosomatid millipedes from India. Most of the genera are reported outside of West Bengal except *Orthomorpha*, *Anoplodesmus*, *Chondromorpha*, *Streptogonopus*, *Oxidus*, *Delarthrum* and *Polydrepanum*. In West Bengal (a state of India), no worker described any genus or species of millipede till today. In my native district Birbhum (West Bengal state), I observed at least five species of millipedes which seems to be new to science. Present paper reports on description of a new genus and species, *Manikidesmus suriensis* of the family Paradoxosomatidae.

## Materials and methods

Millipedes were hand collected from the road side and moist wall during July, 2020. Collections were made in the morning session i. e. 5 a. m. to 10 a. m. at the peak activity period with the most success on overcast day. Specimens were observed in leaf litter and moist wall covered with moss under shady trees.

Hand collected specimens were euthenised by cooling and then preserved in 4% formaldehyde. Specimens were examined using light microscope and sketches were drawn with the help of camera lucida. Photographs were taken from the microscope by camera. Morphological measurements were made with the help of an ocular micrometer.

## Result

### Taxonomy

Class: Diplopoda Blainville in Gervais, 1844 Order: Polydesmida Pocock, 1887

Family: Paradoxosomatidae Daday, 1889

Subfamily: Paradoxosomatinae Daday, 1889

Tribe: Polydrepanini Jeekal, 1968

*Manikidesmus* Bhakat, gen. n.

Type species: *Manikidesmus suriensis* Bhakat, sp. n.

#### Diagnosis

The genus *Manikidesmus* gen. n. is characterized by

1. Reduced paranota, below the midline of the body and in the form of triangle.
2. Unpaired sternal lamella on the 5^th^ sternite, half-circle in shape with blunt end.
3. In male, femur of 2^nd^ segment bears a blunt adenostyle.
4. Prefemur of gonopod bears numerous setae.
5. In the gonopod bearing leg of male, prefemur is swollen with dense setal brush on ventral surface.
6. Gonopod aperture forms two separate opening and lack sternum in male.
7. Gonopod coxa is cylindrical and longer than femur and sparsely setose on dorsal.
8. Acropodite has two branches – solenomere and solenophore.
9. Prefemorite is stout, oval-round and distinctly separated from the acropodite.
10. Gonopod with a distinct postfemoral suture.
11. Trapezoidal spinneret narrowing ventrally, four setae arose from a distinct round setal socket.
12. At the tip of epiproct, a transparent conical cap is present.
13. Hypoproct bears a long seta in the middle.
14. In female, femur of leg is exceptionally longer.
15. Distinct rectangular pleural keel on the anterior segments.
16. Ozopores on segments 5, 7, 9, 10, 12, 13, 15-19.

#### Etymology

The genus is named after Prof. Manik Chandra Mukhopadhyaya of the University of Burdwan as a token appreciation for his contribution to millipede research in West Bengal. The generic epithet is derived from the name “Manik” (used as a noun in the nominative singular) in conjugation with the genus name “desmus” often used as a suffix in Polydesmid generic name. For the purposes of nomenclature, the gender of this genus is male.

#### Remarks

I placed the genus in the tribe Polydrepanini under family Paradoxosomatidae in having the following combination of characters: stout and subcylindrical coxa with a short prefemur, one or more femoral process, post femur is distinctly demarcated by suture, tibiotarsus long cylindrical at its basal end and flattened distally with thin flagelliform solenomere, seminal groove runs diagonally in the femur and almost enclosed by tibiotarsus.

#### Comparison

The new genus is distinguished from other important millipede genera of Indian origin of the family Paradoxosomatidae by several characters. Both *Oxidus* and *Chondromorpha* lack tarsal brush in the leg (Sankaran and Sebastian, 2017; Nguyen et al., 2017) while distinct setal brush is present on tibia and tarsus of the present genus. Moreover, in *Oxidus*, 5^th^ sternum without sternal lamella, while it is well developed in *Manikidesmus* (Nguyen et al., 2017). *Chondromorpha* differs from the present genus by a number of characters like paranota well developed (vs. ill developed), collum with three transverse rows of setae (vs. without setae), pleural keel indistinct (vs. distinct pleural keel), sterna lamella in coxae 4 (vs. in coxae 5) (Sankaran and Sebastian, 2017). In *Chondromorpha*, distal branch of postfemur of gonopod is curved and sickle like while it is straight in *Manikidesmus*. Post femoral spine is absent in the former genus. In the present genus, solenomere is long coiled and flagelliform while it is a curved structure in *Chondromorpha* (Bano and Murthy, 1997). In *Polydrepanum* (Carl, 1932; Attems, 1937; Jeekel, 1968; Golovatch, 1984) and *Orthomorpha* (Bano and Murthy, 1997), the femur of gonopod is long, thin and cylindrical with distinct postfemoral demarcation but in *Manikidesmus*, femur is short, flat and without post femoral demarcation. In *Polydrepanum*, there is no lamina medialis or lamina lateralis but both are present in *Manikidesmus*. In the former genus solenomerite is folded and hidden while it is unfolded and open in the present genus (Bano and Murthy, 1997). In *Anoplodesmus*, tibio-tarsus of gonopod is short, bifid, free and distofemoral process without spine (Nguyen, 2010; Bano and Murthy, 1997) while in the present genus, tibio-tarsus is comparatively long, attached and have a spine in the distofemoral process. Lamina medialis is long straight and possess a curved hook at its distal end in *Manikidesmus* while it is indistinct and without curved hook in *Anoplodesmus*. In *Streptogonopus*, gonopod femorite is distinctly demarcated from postfemoral region and solenomere is twisted with solenophore one or two times (Jeekel, 2004).Compared to *Streptogonopus*, in the present genus there is no demarcation in between femorite and postfemur region and though solenomere is long and coiled but not twisted with solenophore. The present genus is more similar to *Delarthrum* by the presence of setal brush on prefemur, postion of lateral keel on 2^nd^ segment, smooth metazonite, sternite 5^th^ with lamella, pleural keel, short gonopod prefemur. But the later can be distinguished from crus by the combination of following characters: femur of 1^st^ leg of male with adenostyle (vs. 2^nd^ leg with adenostyle), sternite is square in shape (vs. rectangular), tibio-tarsus of gonopod distinctly separated from femur and broad leaf like (vs. not so distinctly separated and comparatively less broad), setal brush on tarsus only (vs. on both tibia and tarsus), lateral keel well developed (vs. ill developed), sterna lamella rectangular in shape (vs. half-circle) (Attems, 1936; Golovatch, 2014). In *Manikidesmus*, prefemur of the gonopod bearing leg of male possesses “setal brush” which is absent in all the Indian genera of Paradoxosomatidae except *Delarthrum*

#### Included species

1. *Manikidesmus suriensis* sp. n. (Fig. 1).

**Fig. 1.**
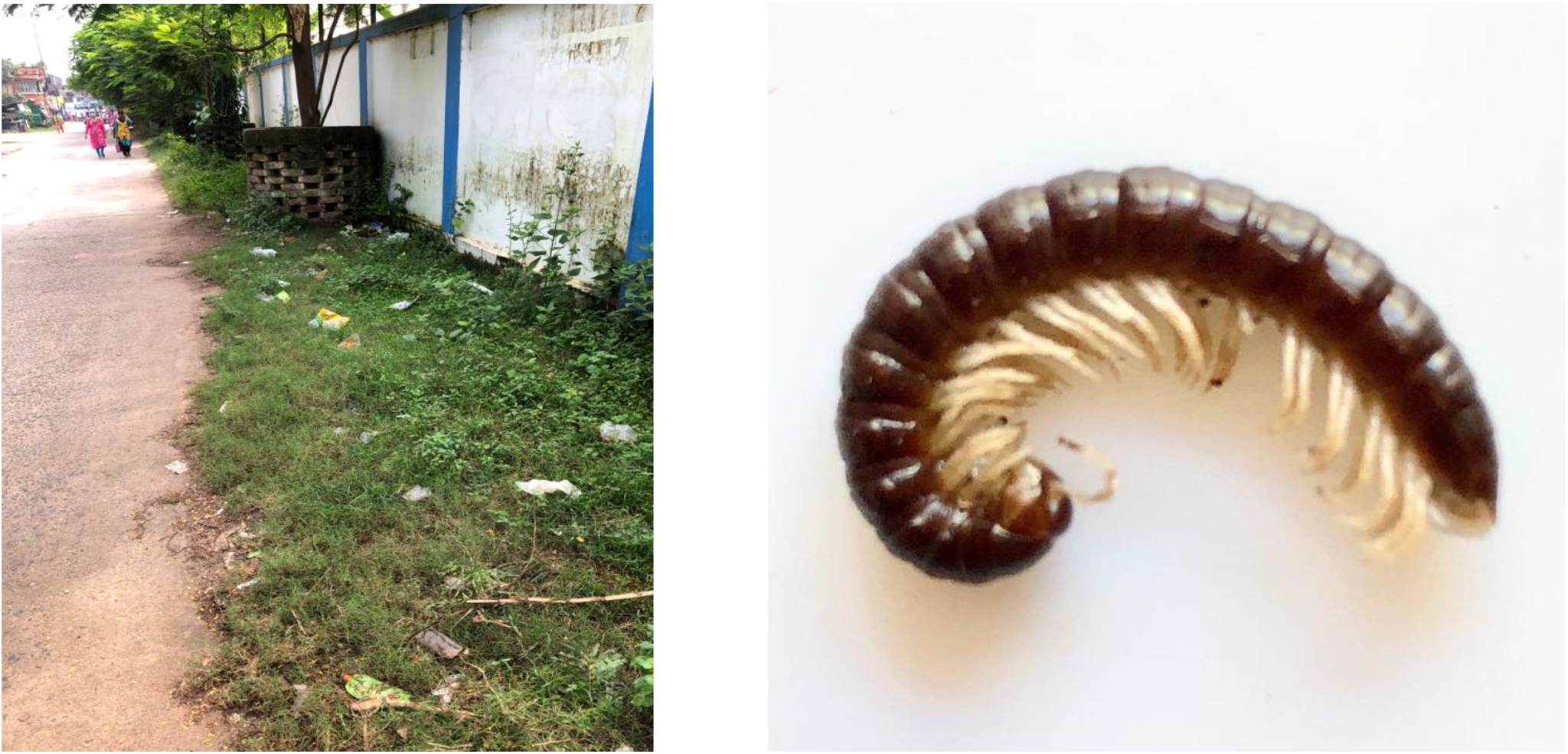
*Manikidesmus suriensis* sp. n. Collection site (left), an adult male (right).

**Fig. 2.**
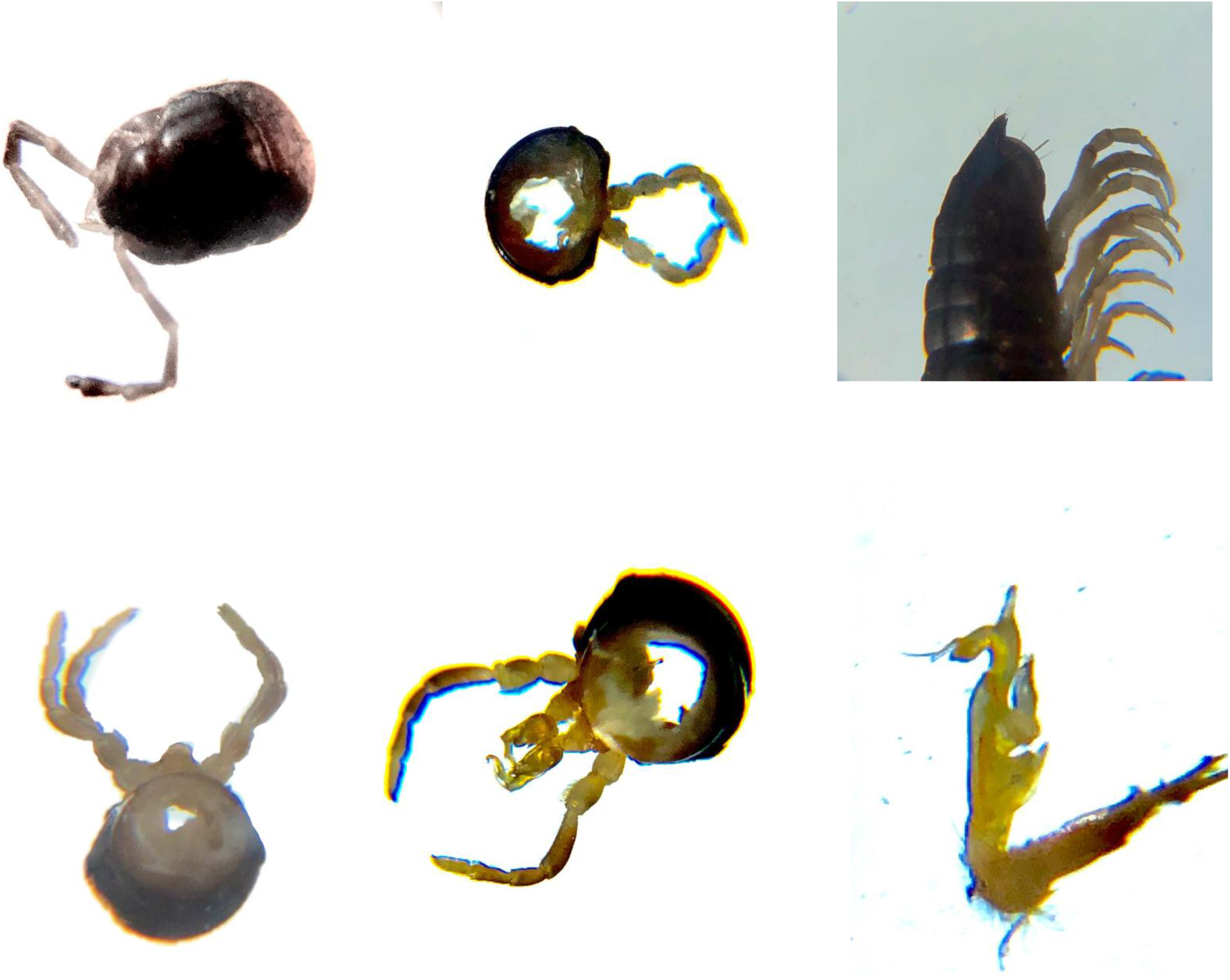
*Manikidesmus suriensis*. 1^st^ row: head with antenna, leg showing adenostyle, telson showing tail 2^nd^ row: segment with sternal lamella, 7^th^ Segment with gonopod in male, gonopod in male. (left to right).

**Fig. 3.**
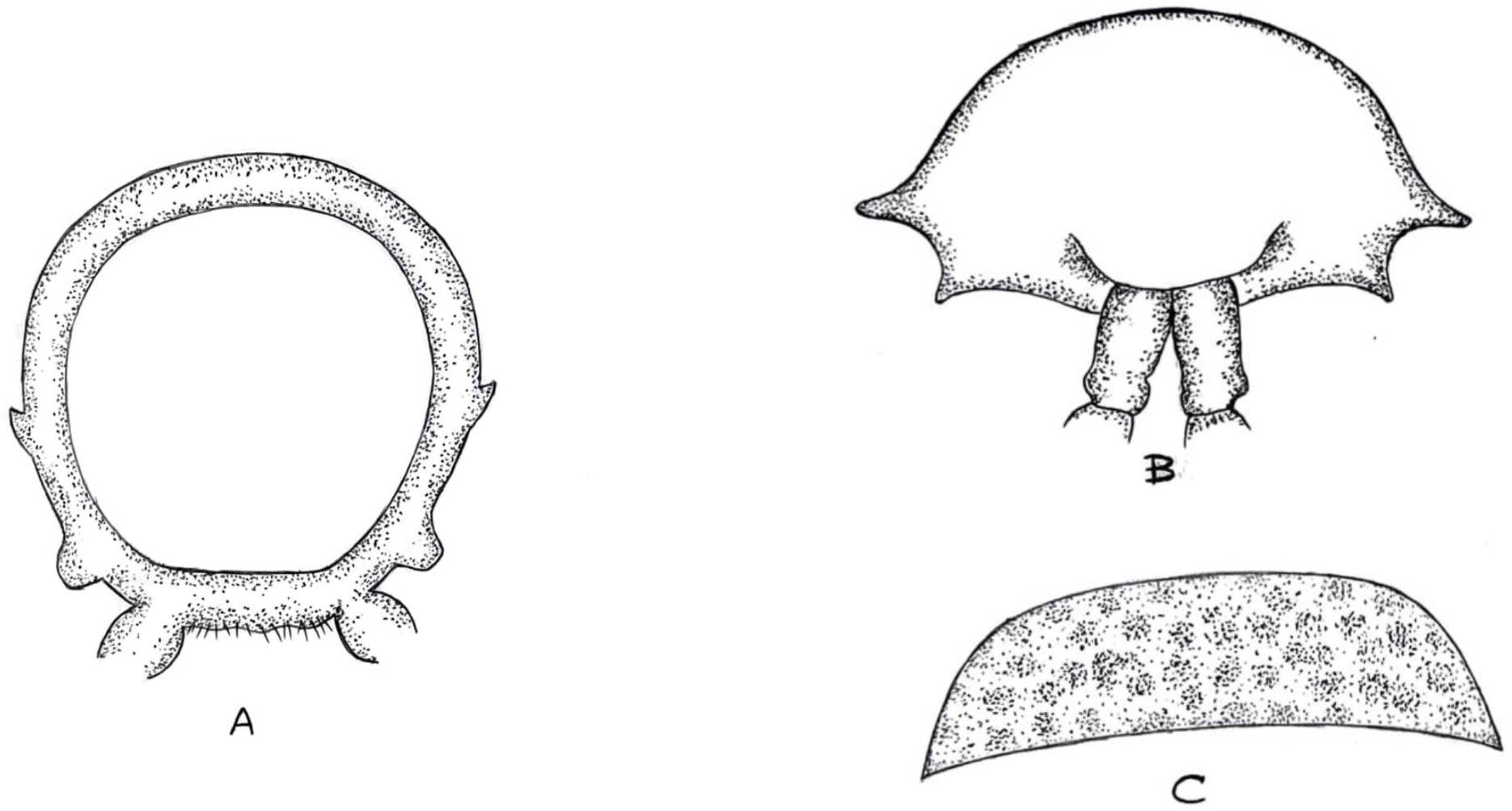
*Manikidesmus suriensis* sp. n. A. Body ring showing paranota and pleural keel, B. 2^nd^ segment, C. Collum.

**Fig. 4.**
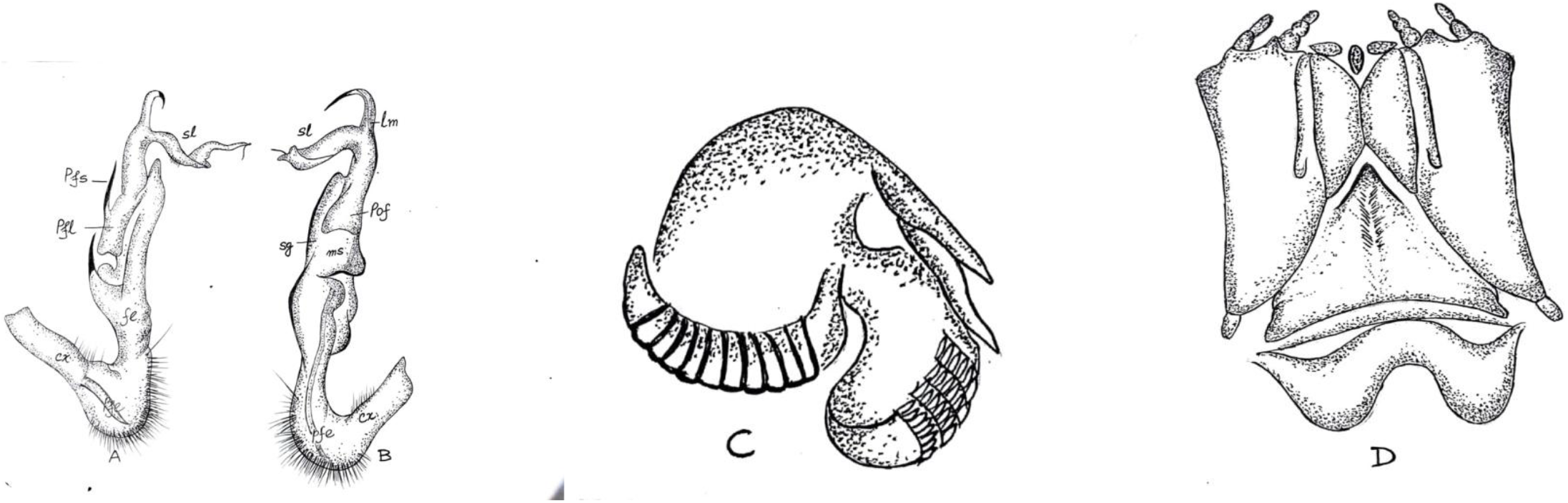
*Manikidesmus suriensis* sp.n. A and B. Gonopod. cx- coxa, pfe- prefemur, fe- femur, msmesal sulcus, pof- post femoral part, sg- seminal groove, pfl- post femoral lamina, pfs- post femoral spine, sl- solenomere, lm- lamina medialis. C. Mandible, D. Gnathochilarium.

**Fig. 5.**
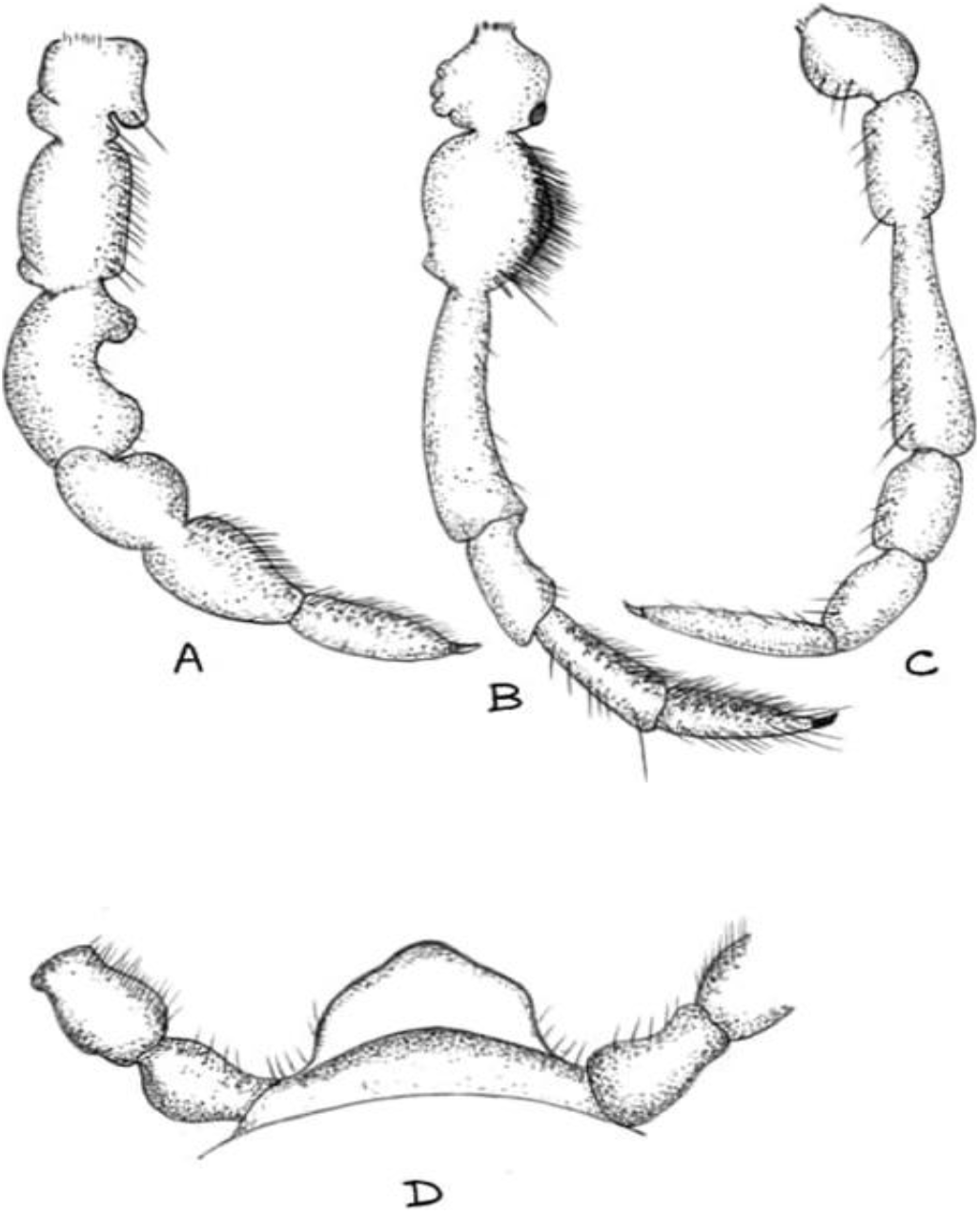
*Manikidesmus suriensis*. A. 2^nd^ leg of male showing adenostyle in femur and setal brush. B. Gonopod bearing leg of male showing trilobed coxa and prefemur with brush. C. Leg of female. D. 5^th^ sternite with sterna lamella.

**Fig. 6.**
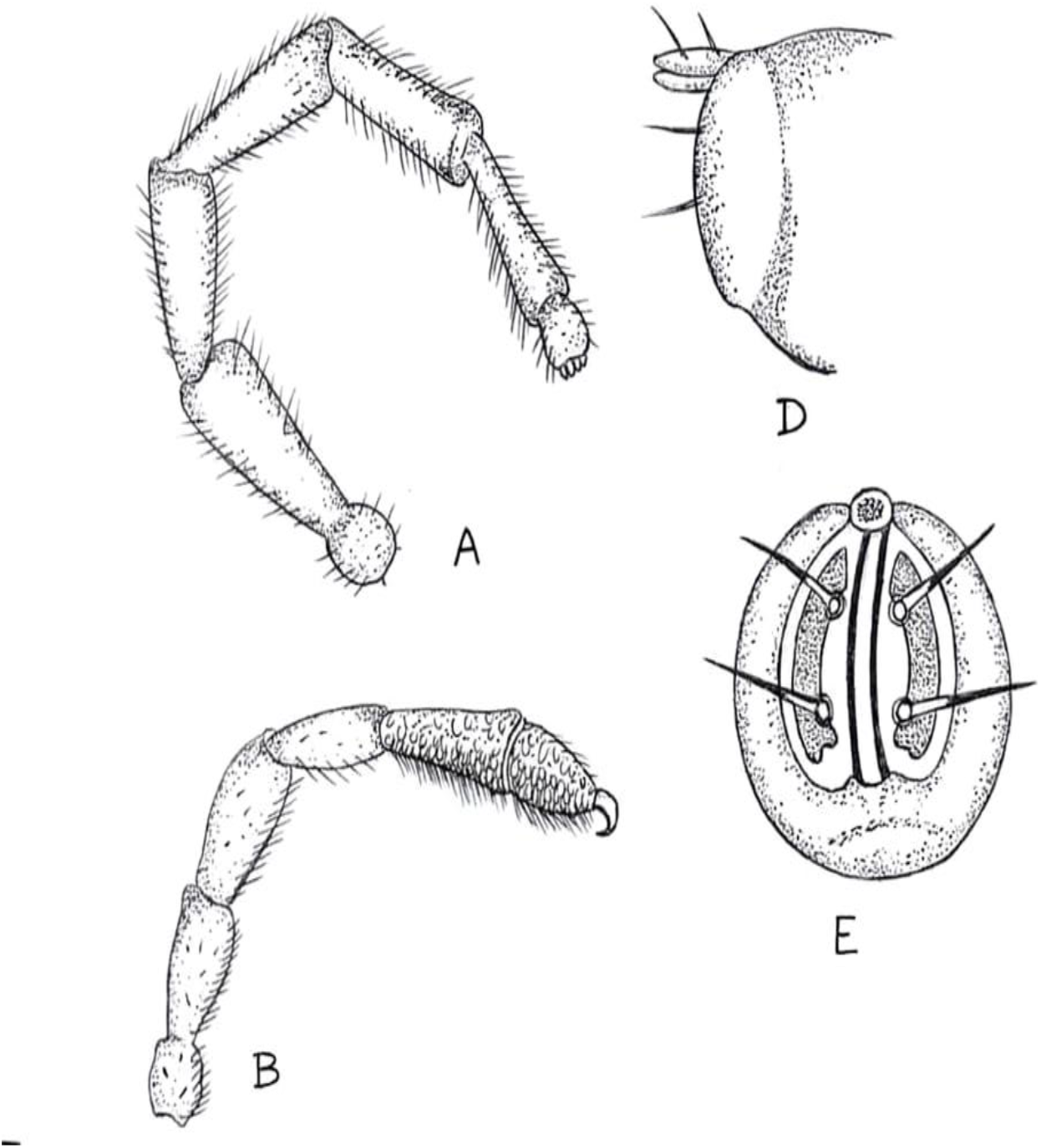
*Manikidesmus suriensis*. A. Antenna, B. Leg of male, D. Telson, E. Anal plate showing paraproct with four long setae and paraproct plate.

#### Type material

Holotype: 2. VII. 2020. Lalkuthipara, Suri (87°32′00′′E, 23°55′00′′N), Birbhum district, West Bengal, India; male, 32.5mm length and 3.15mm width; Zoological Museum, Rampurhat College, Rampurhat-731224, West Bengal, India; Collector: Somnath Bhakat. Paratype: July to September, 2020; same locality; male 31.8-33.2mm (7), female 31.5-34.0mm (4); all other details are same as holotype.

#### Diagnosis

Adult male and female are tanned brown in colour with whitish leg, 20 body rings, reduced paranota, male longer than female, metazona larger than prozona in diameter, prefemur of leg bears setal brush in male, sternal lamella half-circle in shape, blunt adenostyle in the second leg, prefemur of gonopod form knee joint and setose distinctly, solenomere is long, coiled and flagelliform, expanded post femoral lamina, seminal groove is extended beyond post femoral part.

#### Description

Males are longer in size compared to female (length of male 32.5± 0.51mm, female 31.1± 0.62mm) but slender in width (width of male 3.15± 0.21mm, female 3.53± 0.32mm).

Head is with small setae throughout. Collum is subtrapezoidal in dorsal outline, hairless and wider than 2^nd^ segment. Surface of collum mottled. Collum is slightly convex in the middle, more strongly convex towards sides and sides are almost perpendicular. Sternal plate with a few setae and sterna is longer than width (almost four times). In male, 5^th^ sternite bears a distinct unpaired sternal lamella which is protruded like half-circle with blunt tip. Setae on the anterior surface of lamella are small while posterior surface bears a few longer setae. Metaterga is with distinct transverse sulcus. In the body segment, metazona is distinctly larger in diameter than prozona and these two are separated by a deep prominent stricture. In the holotype, prozonite is 2.50mm and metazonite 3.25mm (ratio 1.30). Dorsal surface of metazoan is smooth without any impression. Paranota is reduced to a triangular projection just below the midline of the body. Pleural keel on 6^th^ segment highly developed, rectangular in shape and on 8^th^ segments onwards it is reduced like a conical notch. Like other Paradoxosomatidae, paranotum of the second segment is extended and form triangular projection on both sides (Fig ).

Stipes of gnathochilarium bears a small projection at the posterior part. Mentum bears a triangular ridge at the apex. Each stipes bears a sulcus along the length of lingual plates. In the mandible, inner tooth is slightly longer than the outer tooth and molar plate with 11 distinct ridges (Fig ).

The antenna is long, slender, covered by delicate seta and extended backwards to collum (Fig ). Antenna of female is comparatively shorter than that of male (4.32mm vs. 4.79mm). Antenna in male is 152.06% and in female 122.38% of body diameter. Length of antennomere varies in male and female. In male: 1.18: 3.53: 3.29: 3.76: 3.24: 2.76: 1.0, in female: 1.29: 3.06: 3.06: 3.0: 3.06: 2.47: 1.0. The tip of the antenna bears four conical pegs.

Arrangement of legs: Collum and 2^nd^ ring legless, 3^rd^ and 4^th^ ring – one pair each, 5^th^ to 18^th^ ring – two pairs each, 19^th^ ring and telson legless (in male, the anterior pair of leg is modified to gonopod in the 7^th^ ring). Hence male have 58 and female with 60 legs. Leg of male is significantly longer compared to that of female (4.2mm vs. 3.8mm) (P< 0.05). Leg length in male is 133.33% and in female 106.50% of maximum body width. Length of podomeres also varies in both sexes. In male: 5.0: 5.33: 5.78: 4.0: 5.33: 4.67: 1.0, in female: 3.67: 4.33: 8.22: 2.89: 3.11: 4.67: 1.0. In both sexes femur is the longest part but in female it is exceptionally longer. Compared to female, tibia of male leg is proportionately longer. In male, 2^nd^ leg bear distinct projected outgrowth, adenostyle. Like other paradoxosomatidae, 2^nd^ leg bears gonopore at the distal end of coxa. In male, tibia and tarsus bear bushy setal brush which is more developed in the legs anterior to gonopod bearing segment but in the posterior legs setal brush in tibia is less bushy. Moreover, there are small irregular papillae in the inner surface of both podomeres. In the gonopod bearing leg in male, distal part of coxa bears a round bulge in the anterior while posterior part is trilobed. Moreover, dense brushes of setae are present on the slightly swollen ventral surface of prefemur. (Fig ). In female, leg is simple lacking bulge, or setal brush and the coxa is also simple without any lobe. Ozopores 5, 7, 9, 10, 12, 13 and 15-19.

Coxa of gonopod is longer than femur, stout and cylindrical with a few setae at the dorsal. The prefemur is short (almost half of femur) and round, demarcated from femur by a notch and strongly setose (a long seta at the distal). A slightly curved tube like cannula is present laterally. Femur is long and suberect without torsion with flattened distal part. The mesal side of the femur is grooved along the margin, called seminal groove (sg) which is extended beyond the post femoral part. The solenomerite arises from the distal end of the femur enclosing the seminal groove laterally at its anterior end. Acropodite has two branches – the solenomere and the tibiotarsus or solenophore. Solenomere (sl) is long coiled and flagelliform. Lamina medialis (lm) long, straight and distally with a curved hook like tip. Post femoral lamina (pfl) is expanded and with a long post-femoral spine (pfs). The tibio-tarsus bears one distofemoral process having a spine at the tip (Fig ).

Tail is small, blunt and with a few setae. A transparent conical cap like structure with pointed end at the tip of epiproct. Anal valve is thickly rimmed. Paraproct are evenly convex. Arrangement of spinneret is trapezoidal narrowing ventrally. Each seta is smooth, arises from a distinct round setal socket and longer than non-spinneret setae on epiproct. Hypoproct bears a long seta in the middle (Fig ).

Adult female bears numerous small eggs. A female of 30.00mm length and 3.5 mm width have 710 small whitish round eggs of different sizes of which approximately10% are immature. Sizes of largest and smallest eggs are 0.45 and 0.29mm in diameter respectively.

#### Colouration

The adult millipede is tanned brown in colour with whitish legs. Colour of the antenna is different in both sexes. In female, antennomeres are yellowish white while in male two distal segments are dark brownish. Collum mottled brown. Paranota dark brown while pleural keels are light yellow with brown base. Colour of sternite pale brown. The gonopod in male is yellowish brown in colour.

#### Etymology

The species “*suriensis*” is derived from the name “suri”, type locality of the species.

#### Distribution and habitat

According to the present knowledge, *Manikidesmus suriensis* distributed only at Suri, district Birbhum, W. B., India. The species mostly live in moist soil surface with decomposed leaf litter, moist wall with moss under shady trees. In the early morning they can be seen roaming here and there (Fig. 1). In the same habitat, other species of millipedes are present, the dominant of which is *Anoplodesmus saussurii*.

## Discussion

Prefemur of leg with setal brush is present in the genus *Tylopus* (Nguyen, 2012), *Delarthrum* (Golovatch, 2014) of Paradoxosomatidae (Nguyen, 2012) and in *Victoriombrus* of Dalodesmidae of the order Polydesmida (Mesibov, 2004). Except the present genus and *Delarthrum*, this character is absent in all other genera of Indian Paradoxosomatid millipede.

Golovatch (1984, 1994) and Golovatch and Enghoff (1994) illustrated the morphological details of sterna lamella in male. This structure varies at species and even at genus level and may provide a clue for phylogenetic analysis. Sankaran and Sebastian (2017) provided a key for four Indian species of *Chondromorpha* on the basis of structural variation of sterna lamella in male. Though present observation of this structure is restricted to a single species within a single genus, may be useful for phylogenetic comparison in future.

The arrangement of four setae on the paraproct is designated as “a terminal cluster of four setae” by Hoffman (1982) and referred to as “spinneret” by Mesibov (2006). This structure in the present genus may be used to discriminate genera of Paradoxosomatidae.

Hoffman (1962) classified the gonopod of Strongylsomatidae (Paradoxosomatidae) on the basis of presence or absence of post femoral suture and origin of solenomerite. In *Crusacanthus*, post femoral suture is present and the solenomerite originates from the end of the post femur and thus represent second type.

Pleural keel is most common character in Australian Paradoxosomatid millipede (Australiosomatini) (Jeekel, 1984) and some Indian genus of Paradoxosomatid millipede viz. *Orthomorpha*, *Chondromorpha*, *Streptogonopus*, *Anoplodesmus*, *Harpagomorpha*, *Sundanina*, *Kronopolites* etc. (Attems, 1937; Jeekel, 1980). In some species it is restricted to four or seven somites while in other, it may extend upto 18^th^ somite. In the present genus it is extended to 17^th^ somite though more distinct from 3^rd^ to 7^th^ somite.

Maximum number of mature eggs in female indicates that *Manikidesmus* are semelparous in habit like other Paradoxosomatid millipede (Bhakat et al., 1989).

Based on comparatively large number of specimen and a species of the new genus, the study confirmed that gonopod along with other morphological characters can be used to identify the new genus taxonomically.

## Acknowledgement

I am deeply indebted to my son Dr. Soumendranath Bhakat and my wife for their constant inspiration in this critical period of lock down. I am grateful to my colleagues of Rampurhat College for their cooperation, Mr. Sourav Saha of Satyajit Roy Film and Television Institute, Kolkata for technical help and Dr. Golovatch for his valuable advice and guidance.

